# Landscape functional connectivity for butterflies under different scenarios of land-use, land-cover, and climate change in Australia

**DOI:** 10.1101/2022.02.07.479372

**Authors:** Vishesh L. Diengdoh, Stefania Ondei, Rahil J. Amin, Mark Hunt, Barry W. Brook

## Abstract

Pollinating invertebrates are vital to terrestrial ecosystems but are impacted by anthropogenic habitat loss/fragmentation and climate change. Conserving and improving landscape connectivity is important to offset those threats, yet its assessment for invertebrates is lacking. In this study, we evaluated the functional connectivity between protected areas in Australia for 59 butterfly species, under present conditions and different future scenarios (for 2050 and 2090) of land-use, land-cover, and climate change. Using circuit-theory analysis, we found that functional connectivity under present conditions varies widely between species, even when their estimated geographical ranges are similar. Under future scenarios, functional connectivity is predicted to decrease overall, with negative changes worsening from 2050 to 2090, although a few species are positive exceptions. We have made our results available as spatial datasets to allow comparisons with taxa from other studies and can be used to identify priority areas for conservation in terms of establishing ecological corridors or stepping-stone habitat patches. Our study highlights the importance of considering pollinating invertebrates when seeking holistic conservation and restoration of a landscape’s functional connectivity, underscoring the need to expand and promote protected areas to facilitate functional connectivity under future scenarios of global change.

**Research Data:** The habitat suitability maps and functional connectivity maps are made available as GeoTiff images via Figshare (10.6084/m9.figshare.19130078).

## 1. Introduction

Landscape connectivity is defined as the extent to which the landscape facilitates the movement of organisms between habitat patches. It can be structural or functional; the former is dependent on only the landscape structure, while the latter (which is of focus in this study) also considers species attributes such as habitat preference and dispersal ability (Rudnick et al. 2012; Costanza and Terando 2019). Habitat loss and fragmentation increase barriers between suitable habitats reducing gene flow and the ability of a species to track climate change. Maintaining and restoring landscape connectivity is considered as an important adaptation strategy to reduce the impact of habitat loss and fragmentation and climate change and thereby better conserve biodiversity (Rudnick et al. 2012; Costanza and Terando 2019; Littlefield et al. 2019).

Studies on habitat connectivity (i.e., the degree of functional connectivity between patches of preferred or obligate habitat for individual species) are typically biased towards mammals and birds (Correa Ayram et al. 2016; Dickson et al. 2019). This leaves a gap in the literature for invertebrates—particularly pollinating invertebrates—although there are exceptions (e.g. Filz et al. 2013; Chen 2017; Kirk et al. 2018; Miranda et al. 2021). This gap needs to be filled, as habitat connectivity is important for sustaining pollinator abundance, diversity, and dispersal (Potts et al. 2016). Indeed preserving habitat connectivity is an important strategy for conserving insects (Samways et al. 2020) and its loss can have a negative impact on pollination (Mitchell et al. 2013).

Butterflies make for an ideal functional-connectivity case study because they are a major pollinating taxon thought to be able to transfer pollen over larger distances than other insects (Winfree et al. 2011) and are demonstrably impacted by habitat loss/fragmentation and climate change (Miao et al. 2020; Warren et al. 2021). In some regions, available protected areas are inadequate for butterfly conservation (Chowdhury et al. 2021a) while in other cases, the effectiveness of protected areas for butterflies is predicted to decrease under climate change (Cheng and Bonebrake 2017). Expanding and improving the connectivity of protected areas is part of the Aichi Biodiversity Targets (Target 11; Convention on Biological Diversity 2022). Thus, maintaining and promoting ecological corridors or stepping-stone habitats is critical to facilitate species movement between metapopulations to prevent inbreeding and promote recolonisation after extirpation events (Sands 2018), as well as to facilitate movement in response to climate change (Stewart et al. 2019; Malakoutikhah et al. 2020).

The aim of this study is to assess the landscape-scale functional connectivity for butterflies between protected areas in Australia, under both present conditions and different future scenarios of land-use, land-cover and climate change (for 2050 and 2090). We assessed the connectivity for 59 species of butterflies using the *Circuitscape* Julia package (Anantharaman et al. 2020) which is based on circuit theory (McRae et al. 2008). *Circuitscape* uses circuit theory to predict patterns of movement or dispersal of organisms or genes (McRae et al. 2008) and has been used for studying landscape population genetics and identifying animal movement corridors (Dickson et al. 2019). Circuit theory operates on a continuous layer and considers multiple pathways for movements, making it more flexible than other, simpler methods like the least-cost pathway (McRae et al. 2008). We then considered the conservation implications of our findings for habitat prioritisation.

## 2. Materials and Methods

### 2.1 Butterfly species

Occurrence (presence) data were from the Atlas of Living Australia occurrence downloaded at https://doi.org/10.26197/ala.9028e7dc-2566-44de-8999-dbb36c6685a9 Accessed 15 February 2021. Records were limited to those from 1960 onwards and classified as ‘human observation’. Duplicate records based on latitude and longitude were removed. We used spatial thinning to reduce spatial autocorrelation by removing records closer than a minimum nearest neighbour distance (NMD) using the R package *spThin* (Aiello-Lammens et al. 2015). We followed the method by Amin et al. (2021) to find the optimal NMD for each species separately – adjusting it based on the human activity index. To achieve this, firstly, we stratified Australia into low, medium and high human activity grids of resolution 25 km^2^ using Global Human-Footprint data (Venter et al. 2018). Secondly, we removed all records (i.e., of all species combined) closer than 1.25 km to ensure that data in low-density grids were relatively uncorrelated. Then, for each species, we estimate the threshold number of re-samples to be retained per grid (*h*) using: h= n_i_/N_i_ where *n*_i_ represents the number of samples in low-activity grids and *Ni* is the number of low-activity grids. The threshold values were then applied to calculate the maximum number of re-samples to be retained from medium and high-density grids, using the formula: n_jk_=h × N_jk_, where *n* is the maximum number of re-samples from medium *j* or high *k* density grids, and *N* is the number of medium *j* or high grids *k*. This was repeated 20 times per sampling run and tuned the NMD for high- and medium-activity grids to achieve the maximum number of re-samples (n_*jk*_) for each species.

To enable robust model training and validation, we selected only species with at least 100 unique occurrence records (after removing duplicates and accounting for spatial autocorrelation) for high model performance and estimation of geographical ranges. An exception to this rule was made for the Ptunarra Brown Butterfly (*Oreixencia ptunarra* Couchman 1953) which had 96 records as it is a threatened species (Geyle et al. 2021). We pooled all subspecies into their respective species as this enabled each species to have an adequate number of occurrence records. Consequently, we modelled 59 species in this study, belonging to the Nymphalidae, Lycaenidae, Hesperiidae, Papilionidae families (Supplemental Table A1).

### 2.2 Habitat suitability models

In this study, we used habitat-suitability maps as the ‘resistance layer’ which represents the degree to which the landscape facilitates or blocks the movement of an individual across a given cell. Here, high suitability indicates low resistance (and therefore, a high probability of movement). Habitat suitability models are commonly used to estimate resistance in connectivity models (Correa Ayram et al. 2016), and this approach has the advantage of also providing an opportunity to assess the potential impact of climate change on functional connectivity (Ashrafzadeh et al. 2019; Bonnin et al. 2020; Malakoutikhah et al. 2020).

To model habitat suitability, we selected all 19 bioclimatic variables (version 2.1; https://www.worldclim.org; Fick and Hijmans 2017), elevation (https://www.worldclim.org, and land-use and land-cover change (LULCC; Li et al. 2017) as predictors. The LULCC variable was resampled to match the resolution of the bioclimatic and elevation variables (0.05°, or ~5 km at Australian latitudes) using the *raster* R package (Hijmans 2020). We then reduced the number of predictors by removing highly correlated predictors using a threshold value of 0.7 (Dormann et al. 2013) and implemented using the *findCorrelation* function in *caret* R package (Kuhn 2021). If two variables were highly correlated, then the variable with the largest mean absolute correlation was removed. All non-correlated predictors were used for model fitting for each species (Supplementary Table A1).

The study area for each species was constrained using its estimated kernel geographical range, which was implemented using the *adehabitatHR* R package (Calenge 2006). We assumed that a species can disperse within the entirety of its geographical range.

Pseudo-absences were generated using a random sampling method, with the presence to pseudo-absence ratio set to unity. We used k-fold cross-validation (*k*=10) for algorithm optimisation and implemented using the *caret* R package (Kuhn 2021). The algorithms used in the study include random forest, artificial neural network, k-nearest neighbour, flexible discriminant analysis, and naïve Bayes, as they have different operating mechanisms and so capture a diversity of machine-learning approaches.

Random forest is decisions trees based on bagging (Breiman 2001). Artificial neural network consists of a network of neurons that are considered as the processing units in a strictly feed-forward neural network (Sazli 2006). k-nearest neighbour works under the assumption that similar things exist in proximity (in parameter hyperspace) and thus classifies data most common among its neighbours (Cover and Hart 1967). Flexible discriminant analysis uses multivariate adaptive regression splines to separate the data (Hastie et al. 1994), while naïve Bayes is based on conditional probability (Ren et al. 2009).

Habitat-suitability models were fitted using the trained algorithms, and then ensembled. An ensemble algorithm averages the prediction of structurally different algorithms and has the potential to overcome uncertainty in model selection and improve prediction accuracy by reducing variance and bias (Dormann et al. 2018). As such, we averaged the algorithms using an unweighted method and assessed the goodness of fit using AUC and TSS (Allouche et al. 2006).

To account for the influence of global climate models (GCMs) we selected CanESM5 and MIROC6, which have been used in previous studies in Australia (e.g., Briscoe et al. 2016; Ofori et al. 2017; Morán□Ordóñez et al. 2018) under Shared Socio-economic Pathways (SSP) 7.0, a high-emission scenario (IPCC 2021). For the LULCC variable, we selected A1 (low population growth, sprawling urban expansion, very high economic growth, rapid technological innovation, strong biofuels demand including cellulose-based ethanol) and B1 (low population growth, compact urban expansion, high economic growth, medium technological innovation, low overall energy use, lower demand for biofuels) scenarios which are both oriented globally (Li et al. 2017). Future predictions were made for each species per algorithm per year per climate model per LULCC scenario, and then ensembled (unweighted) per year (2050 and 2090).

### 2.3 Functional connectivity models

Functional connectivity was assessed using the *Circuitscape* package in the Julia Programming Language (Anantharaman et al. 2020). *Circuitscape* calculates connectivity between focal nodes (habitat patches or populations) across a resistance layer (represents the degree to which the landscape impedes the movement of an individual across a given cell) and in analogy to an integrated circuit board in electronics, then calculates the effective resistance and ‘current flow’ which in the ecological interpretation is a measure of net movement probability (McRae et al. 2008). The modelling of functional connectivity results in maps with cumulative current, where the intensity of current is a proxy for a species’ movement at each pixel (Grafius et al. 2017).

In this study, we used the centroids of protected areas as focal nodes (Mukherjee et al. 2021); for this purpose, we only selected protected areas with an average habitat suitability of ≥ 0.7 as this represented a high threshold of suitability. We identified focal nodes for each species individually, under present climate conditions and future climate-change scenarios (for 2050 and 2090).

The resistance layer in our study of butterflies is equal to the sum of the habitat suitability model and Global Human Footprint data (Venter et al. 2018). The Global Human Footprint is a measure of human impact (Venter et al. 2018), where cells with high impact are associated with high resistance, likely to be a function of dispersal limitation rather than habitat suitability, as urban and agricultural areas can reduce the ability of butterflies to move across landscapes and infrastructure such as roads results in significant mortality of butterflies (Chowdhury et al. 2021b). The predicted habitat-suitability data ranges from 0.0 to 1.0 (low to high resistance), while the Global Human Footprint data ranges from 0.0 to 100.0 (high to low). To match the scale and direction of these two data sets, we transformed the habitat suitability data by subtracting the data from 1 and multiplying it by100. The transformed habitat suitability data was added to the Global Human Footprint data and then rounded up to the nearest integer, because *Circuitscape* does not accept non-integer values. Due to the unavailability of future Global Human Footprint data, we were forced to assume that this data is constant under present conditions and future scenarios. The final resistance layer scales from 0 to 200 (lowest to highest possible resistance). Resistance layers were created for each species under present climate conditions and future climate scenarios (for 2050 and 2090).

We predicted the functional connectivity for each species under present conditions and then calculated their goodness of fit (AUC) to assess model accuracy (Jackson et al. 2016). We also calculated the mean cumulative current (with standard error) of the connectivity models. If a pixel facilitates connectivity, then the presence points should on average have higher values than pseudo-absences (Grafius et al. 2017; Rodrigues et al. 2021). We then predicted the functional connectivity for each species under future scenarios (for 2050 and 2090). And assessed the difference in functional connectivity between present conditions and future scenarios (i.e., future minus present functional connectivity model) as well.

## 3. Results

The habitat suitability models for all 59 Australian butterfly species with sufficient data achieved high goodness of fit for both AUC and TSS scores. The highest and lowest AUC scores were 0.99 and 0.94 respectively, while the highest and lowest TSS scores were 0.92 and 0.65 respectively (Supplemental Table A1). The connectivity models also achieved high-to-moderate goodness of fit, with AUC scores ranging between 0.94 and 0.68 (Supplementary Table A1). The mean cumulative current of the presence points was higher than that of the pseudo-absences for functional connectivity models of all species (Supplementary Fig. A1-2).

The circuit-theory results (a proxy for functional connectivity) of all the families except for Papilionidae predicted similar future trends, with mean cumulative current expected to decrease through to 2090 (Fig. 1). However, the results for individual species provided a more nuanced perspective than looking at families and exhibited considerably more variation. For example, *Dispar compacta* (Hesperiidae) has a higher mean cumulative current than *Hypolycaena phorbas* (Lycaenidae) with current predicted to decrease for the former and increase for the latter (Fig. 2b, d), whereas for *Acraea andromacha* (Nymphalidae) and *Graphium choredon* (Papilionidae), future scenarios show similar results to present-day conditions (Fig. 2a, c).

**Fig 1.**
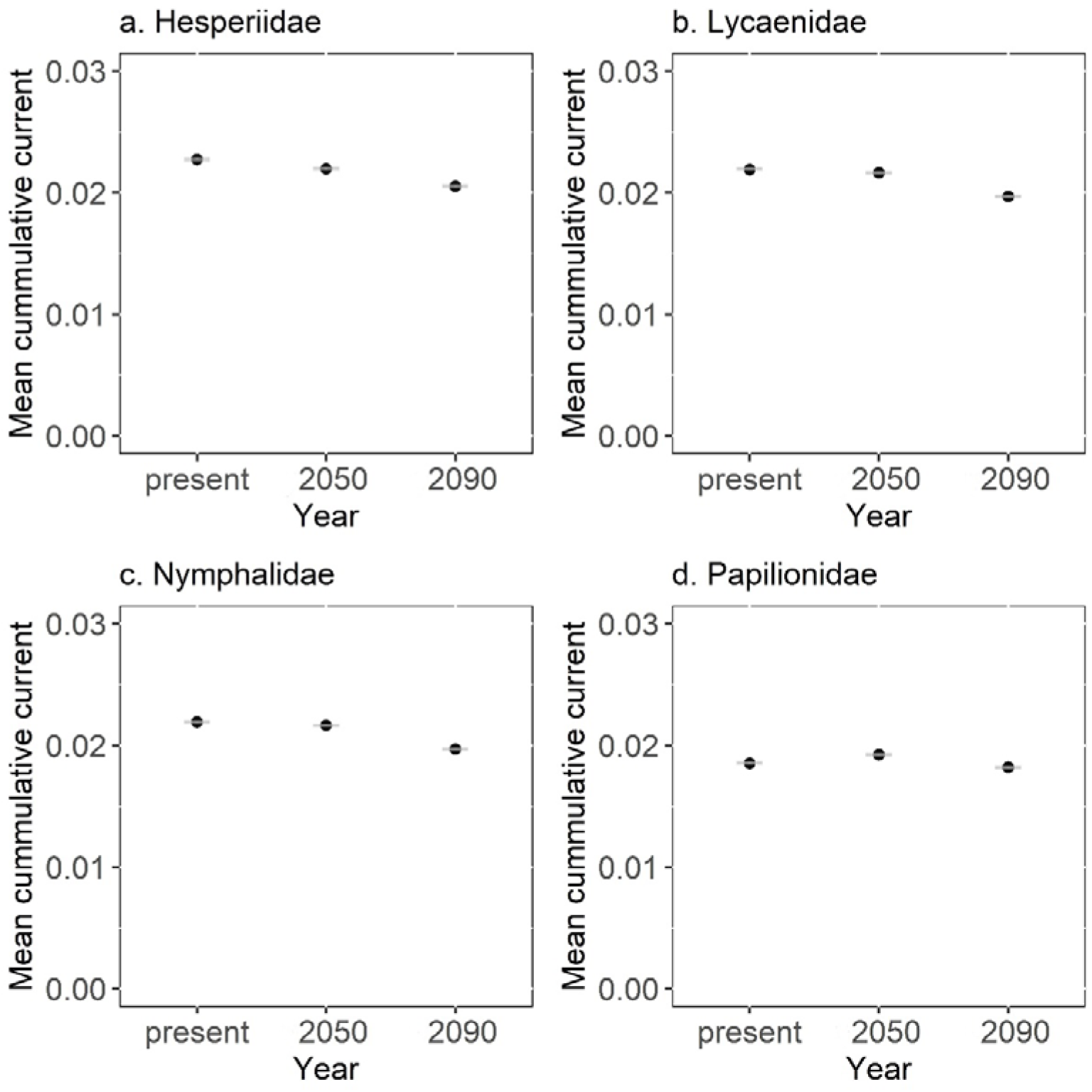
Mean cumulative current with standard error of the families (a) Hesperiidae (b) Lycaenidae, (c) Nymphalidae, and (d) Papilionidae under present conditions and future scenarios of climate change (for the year 2050 and 2090).

**Fig 2.**
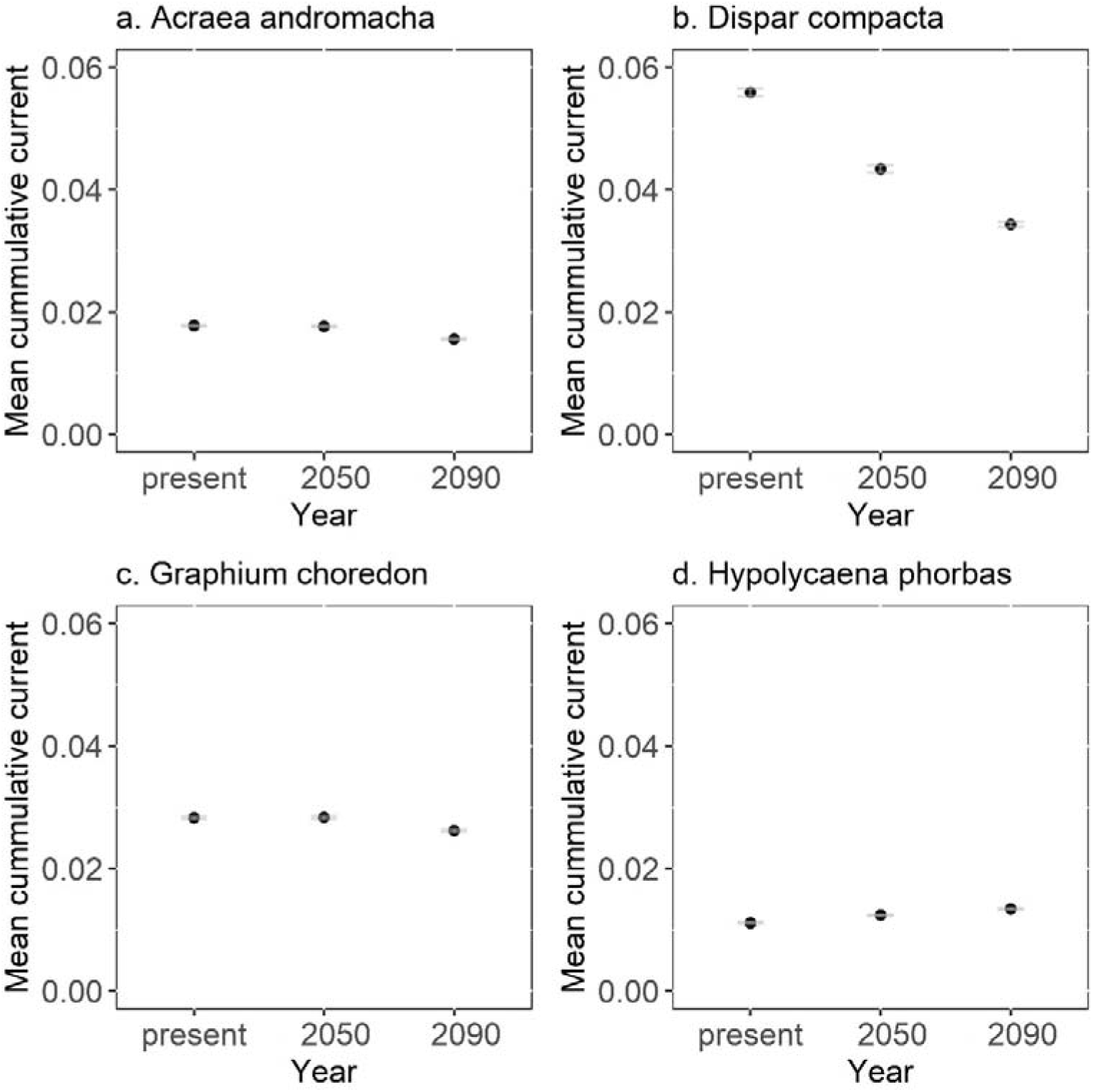
Mean cumulative current with standard error of the species (a) *Acraea andromacha* (b) *Dispar compacta*, (c) *Graphium choredon, and* (d) *Hypolycaena phorbas* under present conditions and future scenarios (the year 2050 and 2090).

Under present conditions, functional connectivity is modelled to vary between species and regionally, including those with similar geographical ranges. For example, *Catochrysops panormus* has higher connectivity along the northern part of Australia than *Graphium eurypylus, Papilio fuscus*, and *Ypthima arctous* (Fig. 3), while *Graphium eurypylus* and *Ypthima arctous* have higher connectivity along the eastern coast than the other species (Fig 3), although those species are found across a similar geographic range.

**Fig 3.**
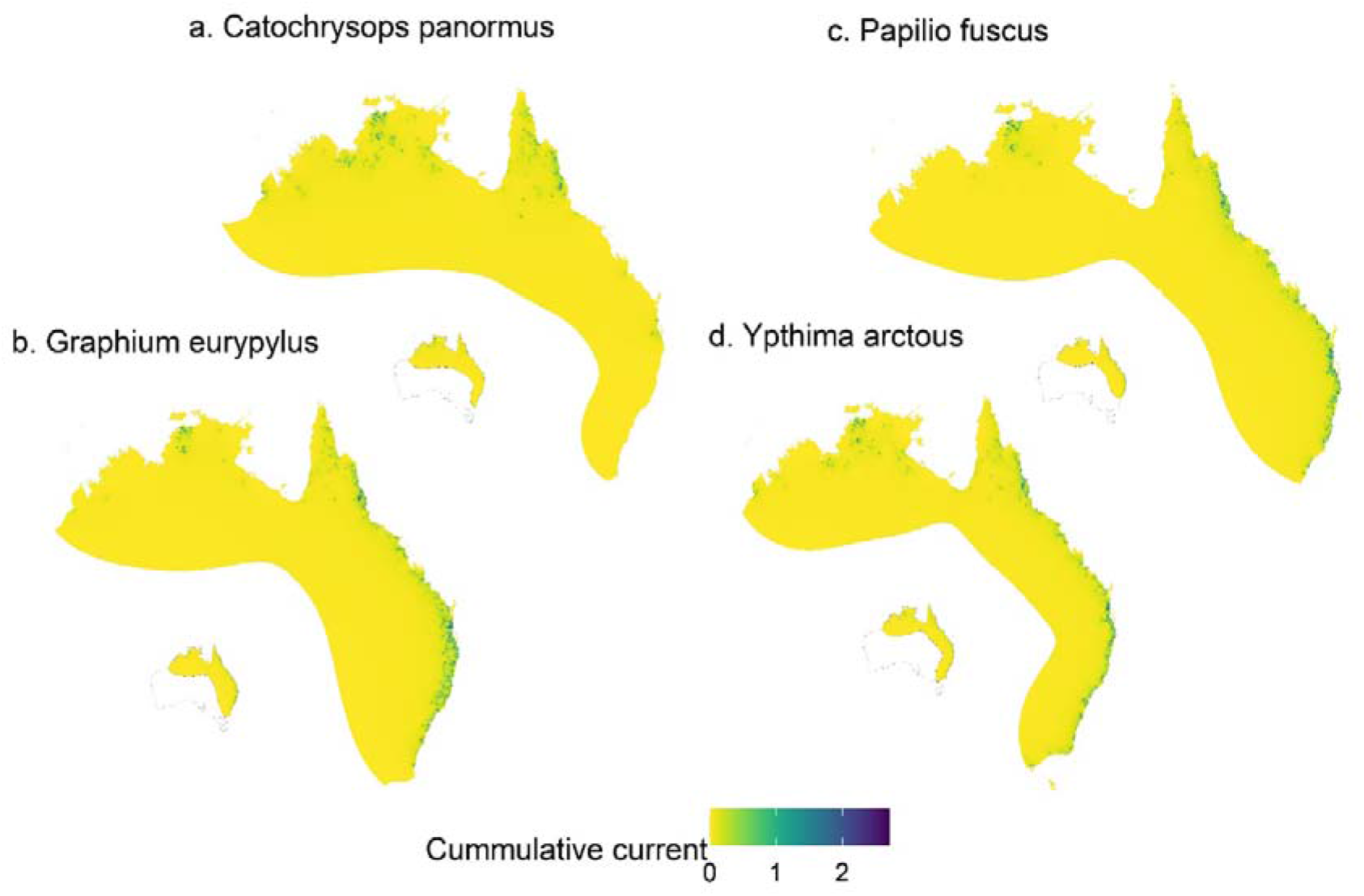
Functional connectivity of the species (a) *Catochrysops panormus* (b) *Graphium eurypylus*, (c) *Papilio fuscus, and* (d) *Ypthima arctous* under present-day conditions.

Most species (43 out of 59) showed consistent, ongoing declines in functional connectivity between the present and 2090 (Fig. 4). For *Arhopala eupolis*, *Candalides erinus*, *Cephrenes augiades*, *Jamides phaseli*, *Nacaduba biocellata*, *Oreixenica ptunarra*, *Papilio fuscus*, *Trapezites symmomus*, and *Vanessa kershawi* (9 out of the 59 species) the percentage of change between future scenarios 2050 and 2090 are similar (Fig 4). While for *Charaxes sempronius*, *Cressida cressida*, *Famegana alsulus*, *Hypolycaena phorbas*, *Junonia hedonia*, *Pelopidas lyelli*, and *Zizeeria karsandra* (7 out of the 59 species) the percentage of positive changes is higher in the year 2090 than in 2050 (Fig 4).

**Fig 4.**
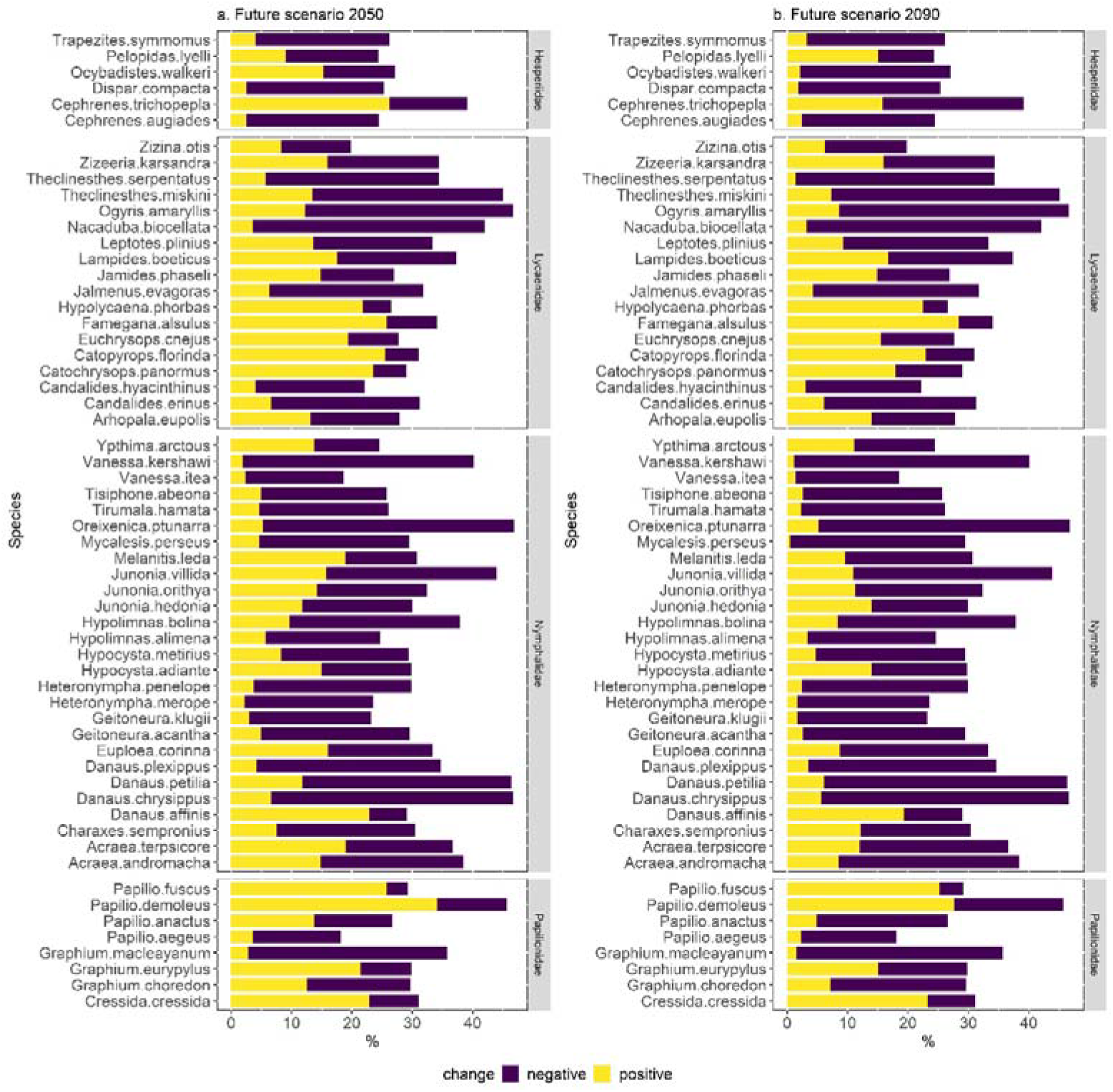
Percentage of negative and positive changes in functional connectivity for each Australian butterfly species, grouped by family, under future scenarios (a) 2050 and (b) 2090.

Although the percentage of change between future scenarios 2050 and 2090 are similar, for the *Oreixenica ptunarra* (Fig. 4) functional connectivity is still predicted to decrease, particularly along the north-west and eastern parts of its range (Fig. 5a-c). And while the percentage of positive changes is higher in the year 2090 than 2050 for *Pelopidas lyelli* and *Famegana alsulus* (Fig 4) functional connectivity is predicted to increase along the southern part of the range for *Pelopidas lyelli* (Fig. 5d-f) and along the north-eastern part of the range for *Famegana alsulus* (Fig. 5g-i)

**Fig 5.**
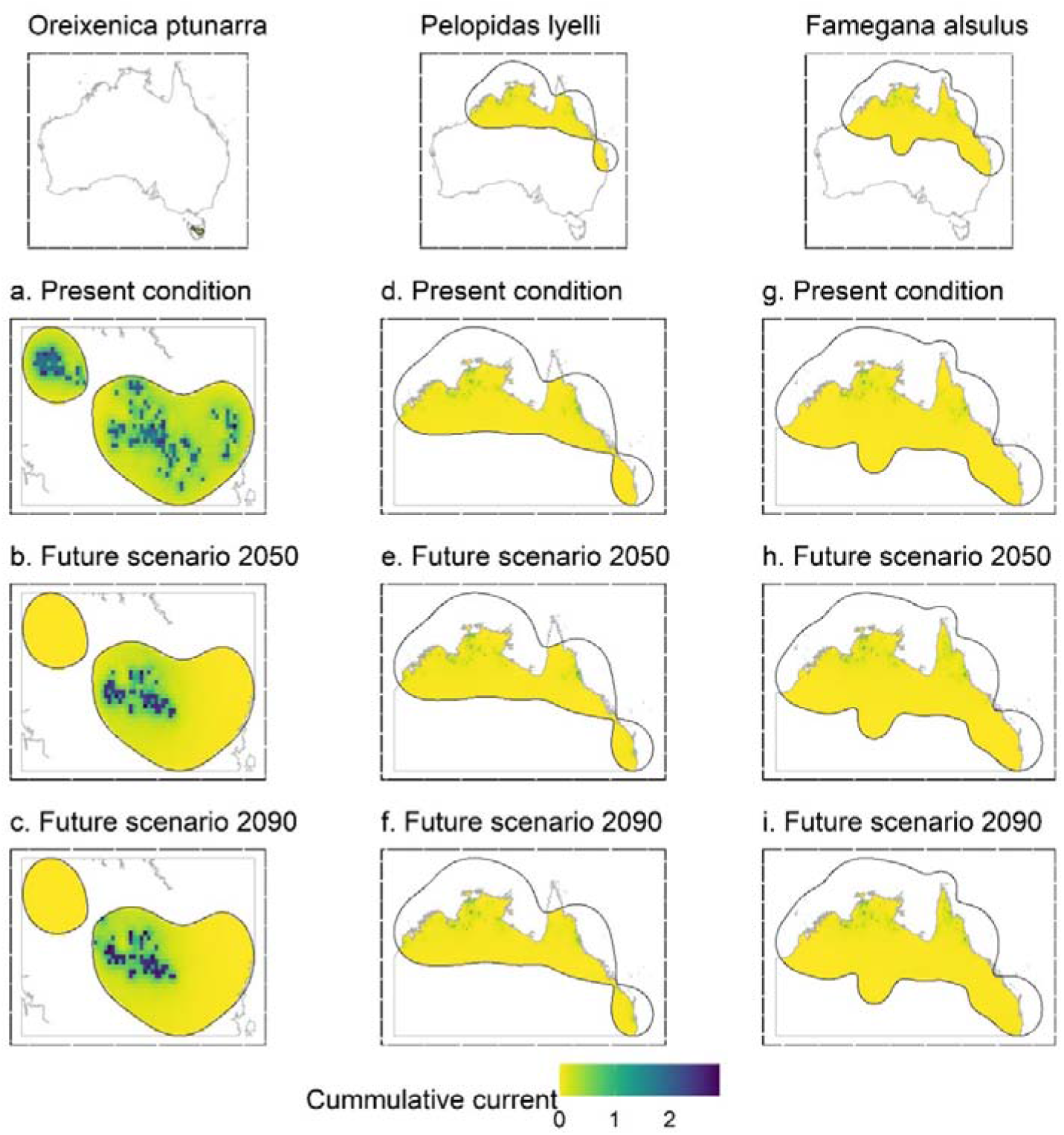
Functional connectivity of the species (a-c) *Oreixenica ptunarra*, (d-f) *Pelopidas lyelli*, and (g-i) *Famegana alsulus* under present conditions and future scenarios (for 2050 and 2090).

Changes in functional connectivity were predicted to occur both along the periphery of a species’ geographical range and within the core. For example, functional connectivity was predicted to be lost along the north-east range for *Geitoneura klugii* and *Heteronympha Penelope* (Fig. 6) as well as within the core of their ranges. In some cases, functional connectivity is also predicted to increase along the edges, for example, there is an increase along the south-west of the island of Tasmania for the *Geitoneura klugii* (Fig. 6a, b).

**Fig 6.**
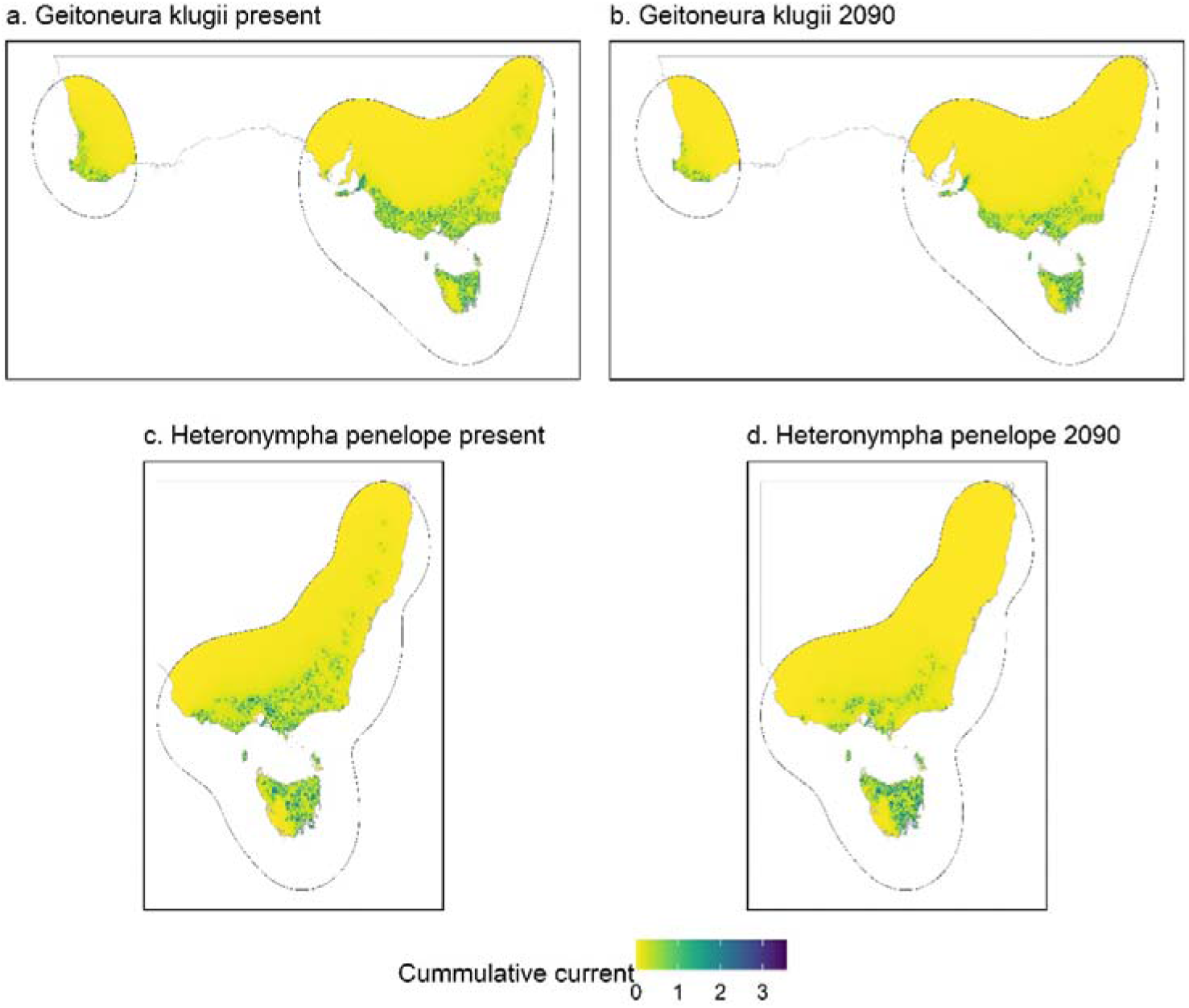
Functional connectivity of *Geitoneura klugii* (a-b) and *Heteronympha penelope* (c-d) under present conditions and future scenarios.

## 4. Discussion

Butterflies are an ecologically important and conspicuous pollinating taxon that is threatened by habitat loss/fragmentation and climate change (Miao et al. 2020; Warren et al. 2021). These threats can be mitigated by conserving and promoting functional connectivity, making it crucial that ecologists seek to identify such areas. Overall, our analysis predicts that functional connectivity will show an overall decrease, with most butterfly species experiencing a higher percentage of negative changes than positive; a trend that worsens over time. Below we highlight how modelling can assist in the decision making of where ecological corridors and stepping-stone habitats should be prioritised.

Under present conditions, the mean cumulative current is overall predicted to be higher along areas with core habitats or focal nodes (i.e., areas with high habitat suitability); a finding common to studies of other taxa such as African elephants (Roever et al. 2013), birds (Grafius et al. 2017), and ungulates (Malakoutikhah et al. 2020). The differences and similarities in functional connectivity between butterfly species with similar (estimated) ranges can be attributed, in large part, to their habitat preferences. For example, *C. panormus* (functional connectivity predicted to decrease; Fig 4) occur in open eucalyptus forest and savannah woodland, whereas *G. eurypylus* (functional connectivity predicted to decrease; Fig 4) occur in monsoon forest, rainforest, and even urban gardens (Braby 2000). As another relevant contrast of species with similar-sized geographic ranges but different responses are, *P. fuscus*, (functional connectivity predicted to remain similar between 2050 and 2090; Fig 4) being found in coastal and subcoastal lowlands rainforest and monsoon forest, compared to *Y. arctous* (functional connectivity predicted to decrease; Fig 4), which prefers coastal and subcoastal woodlands and open forest (Braby 2000).

Overall, we predicted the functional connectivity of most butterfly species in Australia to decrease over the coming decades, albeit with a few exceptions. Our predictions are similar to studies that predicted several non-butterfly taxa such as the Sichuan snub nosed monkey (Zhang et al. 2019b), ungulates (Malakoutikhah et al. 2020; Liang et al. 2021), and the Himalayan brown bear (Mukherjee et al. 2021) to experience a future decrease in functional connectivity due to climate change in different parts of the world. To our knowledge, this is the first attempt to predict the combined impacts of land-use, land-cover, and climate change on the functional connectivity of butterflies.

The mean cumulative circuit-theory ‘current’ is overall predicted to be highest along areas with the best habitat suitability for a given species. The predicted decrease in functional connectivity for most species is expected because climate change is predicted to change the geographic distributions of butterflies (Adhikari et al. 2020; Minachilis et al. 2021). Overall, most of the changes are predicted along the edges of a species range, because populations along boundaries are generally inhabiting the limits of their physiological tolerances compared to those at the core, leaving them more vulnerable to climate change (Parmesan et al. 2000). These changes could also be due to land-use and land-cover change which is observed (Zhang et al. 2019a; Wang et al. 2020) and predicted (Li et al. 2017) to result in loss of forest areas which could have negative impacts on species depending on such habitats. Overall land-use and land-cover change have a negative impact on biodiversity in Australia including on butterflies (Thackway 2018; Davidson et al. 2021; Kutt et al. 2021).

Given the continental scale of the study area and the number of species assessed, there were a few limitations to the study. Spatial scale can influence functional connectivity models (Laliberté and St-Laurent 2020) and while 1-km spatial resolution predictors are available, the extent of the study area and the high computational requirements forced us to use a coarser resolution of 5 km. The accuracy of the habitat suitability models can be influenced by several factors, including the temporal equilibrium (or lack thereof) between data points (species observations) and the geophysical and landscape predictors, as well as the interaction between the spatial scale of the predictors and attributes of the species (Dormann 2007). Validating functional connectivity models is a challenging process (Laliberté and St-Laurent 2020), with suggested methods including field observations by scientists or automated field recorders (e.g., camera traps, acoustic recorders), along with accurate GPS data (Grafius et al. 2017; Finch et al. 2020; Laliberté and St-Laurent 2020). However, in this study, we used citizen science data to build and validate the models, because of the scale of the study area and the number of species studied, which has the advantage of volume, but the constraint of a lower-level of precision and quality control.

## 5. Conclusion

Butterflies are an important pollinating group, but the functional connectivity for several species are predicted to decrease across Australia in the coming decade due to the combined impacts of land-use, land-cover and climate change. Conservation efforts are being made to improve ecological corridors and stepping-stone habitat-restoration programs to promote functional connectivity, in some cases these efforts include invertebrates such as bees (e.g., Miranda et al. 2021) in other cases the focus is only on vertebrate taxa (e.g., Jones et al. 2021). We advocate conservation efforts should include butterflies and other pollinating taxa as well. The availability of our results as a spatial dataset, along with analogous findings from other taxa, will assist in identifying priority conservation areas. Future studies on butterflies should consider (1) collecting dispersal data, to build better connectivity models given that radio telemetry for butterflies is now becoming a logistically viable option (Wang et al. 2019), (2) improving the resistance layer by including spatial data that contains food plants that caterpillars feed upon and butterflies pollinate (Kass et al. 2020), and (3) focussing on species most threatened (Geyle et al. 2021), to develop more targeted, species-specific conservation efforts.

## Acknowledgements

This work was supported by the Australian Research Council [grant number FL160100101].

