## Supplementary Material for "Landscape functional connectivity for butterflies under different scenarios of land-use, land-cover, and climate change in Australia"

**Supplementary Information**

**Table A1.** List of species (with the number of presence points-n), their predictors used for model fitting and the accuracy scores of the habitat suitability models and functional connectivity models. Please note: with the exception of Oreixenica ptunarra which was stated to have less than 100 points in the main text the other species have less than 100 presence points due to the removal of NA’s.

| **Family** | **Species** | **n** | **Predictors** | **Habitat suitability model** | | **Functional connectivity model** |
| --- | --- | --- | --- | --- | --- | --- |
|  |  |  |  | **AUC** | **TSS** | **AUC** |
| Nymphalidae | Acraea andromacha | 331 | lulc + bio9+ bio14+ bio18+ dem | 0.958 | 0.712 | 0.858 |
| Nymphalidae | Acraea terpsicore | 153 | lulc+ bio8+ bio15+ bio16+ bio19+ dem | 0.969 | 0.732 | 0.853 |
| Lycaenidae | Arhopala eupolis | 124 | lulc+ bio3+ bio8+ bio12+ bio15+ bio18+ dem | 0.980 | 0.838 | 0.690 |
| Lycaenidae | Candalides erinus | 194 | lulc+ bio8+ bio15+ bio18+ dem | 0.967 | 0.711 | 0.722 |
| Lycaenidae | Candalides hyacinthinus | 121 | lulc+ bio3+ bio4+ bio6+ bio8+ bio9+ bio15+ bio18+ bio19+ dem | 0.985 | 0.801 | 0.887 |
| Lycaenidae | Catochrysops panormus | 104 | lulc+ bio2+ bio3+ bio17+ dem | 0.967 | 0.740 | 0.863 |
| Lycaenidae | Catopyrops florinda | 146 | lulc+ bio17+ bio18+ dem | 0.964 | 0.767 | 0.731 |
| Hesperiidae | Cephrenes augiades | 108 | lulc+ bio2+ bio8+ bio9+ bio17+ bio18+ dem | 0.997 | 0.925 | 0.904 |
| Hesperiidae | Cephrenes trichopepla | 119 | lulc+ bio2+ bio9+ bio16+ bio19+ dem | 0.980 | 0.815 | 0.783 |
| Nymphalidae | Charaxes sempronius | 175 | lulc+ bio5+ bio18+ dem | 0.932 | 0.673 | 0.784 |
| Papilionidae | Cressida cressida | 224 | lulc + bio17+ bio18+ dem | 0.973 | 0.75 | 0.833 |
| Nymphalidae | Danaus affinis | 246 | lulc+ bio14+ bio18 + dem | 0.983 | 0.780 | 0.656 |
| Nymphalidae | Danaus chrysippus | 117 | lulc+ bio2+ bio3+ bio9+ bio15+ bio18+ bio19+ dem | 0.964 | 0.709 | 0.819 |
| Nymphalidae | Danaus petilia | 646 | lulc+ bio2+ bio8+ bio9+ bio16+ dem | 0.941 | 0.651 | 0.767 |
| Nymphalidae | Danaus plexippus | 473 | lulc+ bio3+ bio8+ bio9+ bio14+ bio15+ bio18+ bio19+ dem | 0.988 | 0.839 | 0.925 |
| Hesperiidae | Dispar compacta | 98 | lulc+ bio1+ bio3+ bio6+ bio7+ bio8+ bio9+ bio13+ bio14+ bio15+ bio19+ dem | 0.981 | 0.785 | 0.865 |
| Lycaenidae | Euchrysops cnejus | 172 | lulc+ bio14+ bio18+ dem | 0.970 | 0.656 | 0.785 |
| Nymphalidae | Euploea corinna | 423 | lulc+ bio9+ bio17+ bio18+ dem | 0.974 | 0.716 | 0.768 |
| Lycaenidae | Famegana alsulus | 215 | lulc+ bio2+ bio3+ bio8+ bio15+ bio19+ dem | 0.972 | 0.753 | 0.712 |
| Nymphalidae | Geitoneura acantha | 184 | lulc+ bio3+ bio4+ bio8+ bio9+ bio14+ bio15+ bio18+ bio19+ dem | 0.970 | 0.787 | 0.842 |
| Nymphalidae | Geitoneura klugii | 224 | lulc+ bio3+ bio4+ bio6+ bio8+ bio9+ bio15+ bio18+ bio19+ dem | 0.976 | 0.785 | 0.889 |
| Papilionidae | Graphium choredon | 168 | lulc+ bio2+ bio3+ bio8+ bio9+ bio19+ dem | 0.995 | 0.910 | 0.946 |
| Papilionidae | Graphium eurypylus | 129 | lulc+ bio3+ bio17+ bio18+ dem | 0.975 | 0.759 | 0.786 |
| Papilionidae | Graphium macleayanum | 185 | lulc+ bio4+ bio8+ bio9+ bio15+ bio18+ dem | 0.994 | 0.843 | 0.895 |
| Nymphalidae | Heteronympha merope | 907 | lulc+ bio3+ bio4+ bio6+ bio8+ bio9+ bio15+ bio18+ bio19+ dem | 0.982 | 0.751 | 0.889 |
| Nymphalidae | Heteronympha penelope | 114 | lulc+ bio3+ bio4+ bio6+ bio8+ bio9+ bio13+ bio15+ bio18+ dem | 0.999 | 0.868 | 0.860 |
| Nymphalidae | Hypocysta adiante | 317 | lulc+ bio14+ bio18+ dem | 0.947 | 0.690 | 0.737 |
| Nymphalidae | Hypocysta metirius | 213 | lulc + bio3+ bio7+ bio8+ bio9+ bio14+ dem | 0.994 | 0.920 | 0.929 |
| Nymphalidae | Hypolimnas alimena | 94 | lulc+ bio3+ bio7+ bio8+ bio19+ dem | 0.973 | 0.763 | 0.694 |
| Nymphalidae | Hypolimnas bolina | 517 | lulc + bio3+ bio8+ bio17+ bio18+ dem | 0.965 | 0.717 | 0.837 |
| Lycaenidae | Hypolycaena phorbas | 81 | lulc+ bio2+ bio3+ bio14+ bio18+ dem | 0.997 | 0.913 | 0.730 |
| Lycaenidae | Jalmenus evagoras | 118 | lulc+ bio1+ bio3+ bio4+ bio9+ bio14+ bio15+ dem | 0.986 | 0.686 | 0.866 |
| Lycaenidae | Jamides phaseli | 97 | lulc+ bio9+ bio14+ bio18+ dem | 0.985 | 0.752 | 0.753 |
| Nymphalidae | Junonia hedonia | 179 | lulc+ bio4+ bio8+ bio9+ bio15+ bio18+ dem | 0.961 | 0.726 | 0.737 |
| Nymphalidae | Junonia orithya | 370 | lulc + bio3+ bio17+ bio18+ dem | 0.964 | 0.740 | 0.736 |
| Nymphalidae | Junonia villida | 1071 | lulc+ bio8+ bio9+ bio12+ bio17+ dem | 0.949 | 0.716 | 0.822 |
| Lycaenidae | Lampides boeticus | 139 | lulc+ bio2+ bio8+ bio9+ bio14+ bio18+ dem | 0.944 | 0.667 | 0.747 |
| Lycaenidae | Leptotes plinius | 100 | lulc+ bio3+ bio5+ bio8+ bio9+ bio12+ dem | 0.976 | 0.79 | 0.904 |
| Nymphalidae | Melanitis leda | 326 | lulc+ bio3+ bio17+ bio18+ dem | 0.983 | 0.828 | 0.866 |
| Nymphalidae | Mycalesis perseus | 109 | lulc+ bio3+ bio7+ bio8+ bio15+ bio19+ dem | 0.989 | 0.834 | 0.721 |
| Lycaenidae | Nacaduba biocellata | 191 | lulc+ bio2+ bio8+ bio9+ bio12+ bio17+ dem | 0.960 | 0.769 | 0.695 |
| Hesperiidae | Ocybadistes walkeri | 269 | lulc + bio8+ bio9+ bio12+ bio14+ dem | 0.980 | 0.821 | 0.891 |
| Lycaenidae | Ogyris amaryllis | 112 | lulc+ bio2+ bio3+ bio8+ bio9+ bio17+ bio18+ bio19+ dem | 0.973 | 0.794 | 0.706 |
| Nymphalidae | Oreixenica ptunarra | 96 | lulc+ bio3+ bio4+ bio6+ bio9+ bio18+ dem | 0.991 | 0.645 | 0.755 |
| Papilionidae | Papilio aegeus | 413 | lulc+ bio3+ bio8+ bio9+ bio12+ bio14+ bio15+ dem | 0.977 | 0.808 | 0.903 |
| Papilionidae | Papilio anactus | 221 | lulc+ bio2+ bio3+ bio8+ bio9+ bio14+ bio15+ bio18+ bio19+ dem | 0.972 | 0.775 | 0.912 |
| Papilionidae | Papilio demoleus | 253 | lulc+ bio2+ bio4+ bio8 + bio9+ dem | 0.946 | 0.687 | 0.748 |
| Papilionidae | Papilio fuscus | 129 | lulc+ bio15+ bio18+ bio19+ dem | 0.983 | 0.868 | 0.765 |
| Hesperiidae | Pelopidas lyelli | 142 | lulc+ bio3+ bio13+ bio17+ bio18+ dem | 0.963 | 0.676 | 0.742 |
| Lycaenidae | Theclinesthes miskini | 199 | lulc+ bio8+ bio9+ bio12+ dem | 0.971 | 0.743 | 0.784 |
| Lycaenidae | Theclinesthes serpentatus | 139 | lulc+ bio8+ bio9+ bio12+ dem | 0.950 | 0.695 | 0.847 |
| Nymphalidae | Tirumala hamata | 301 | lulc + bio3+ bio5 + bio18+ dem | 0.977 | 0.767 | 0.878 |
| Nymphalidae | Tisiphone abeona | 242 | lulc + bio3+ bio7+ bio8+ bio9+ bio13+ bio14+ bio15+ bio19+ dem | 0.992 | 0.867 | 0.815 |
| Hesperiidae | Trapezites symmomus | 156 | lulc+ bio3+ bio7+ bio8+ bio9+ bio14+ dem | 0.982 | 0.826 | 0.892 |
| Nymphalidae | Vanessa itea | 499 | lulc+ bio3+ bio4+ bio8+ bio9+ bio15+ bio18+ dem | 0.968 | 0.775 | 0.885 |
| Nymphalidae | Vanessa kershawi | 878 | lulc+ bio3+ bio4+ bio6+ bio8+ bio9+ bio13+ bio15+ dem | 0.968 | 0.767 | 0.893 |
| Nymphalidae | Ypthima arctous | 203 | lulc+ bio17+ bio18+ dem | 0.962 | 0.714 | 0.686 |
| Lycaenidae | Zizeeria karsandra | 194 | lulc+ bio2+ bio17+ bio18+ dem | 0.991 | 0.845 | 0.745 |
| Lycaenidae | Zizina otis | 189 | lulc+ bio2+ bio3+ bio8+ bio9+ bio17+ bio18+ bio19+ dem | 0.985 | 0.825 | 0.900 |

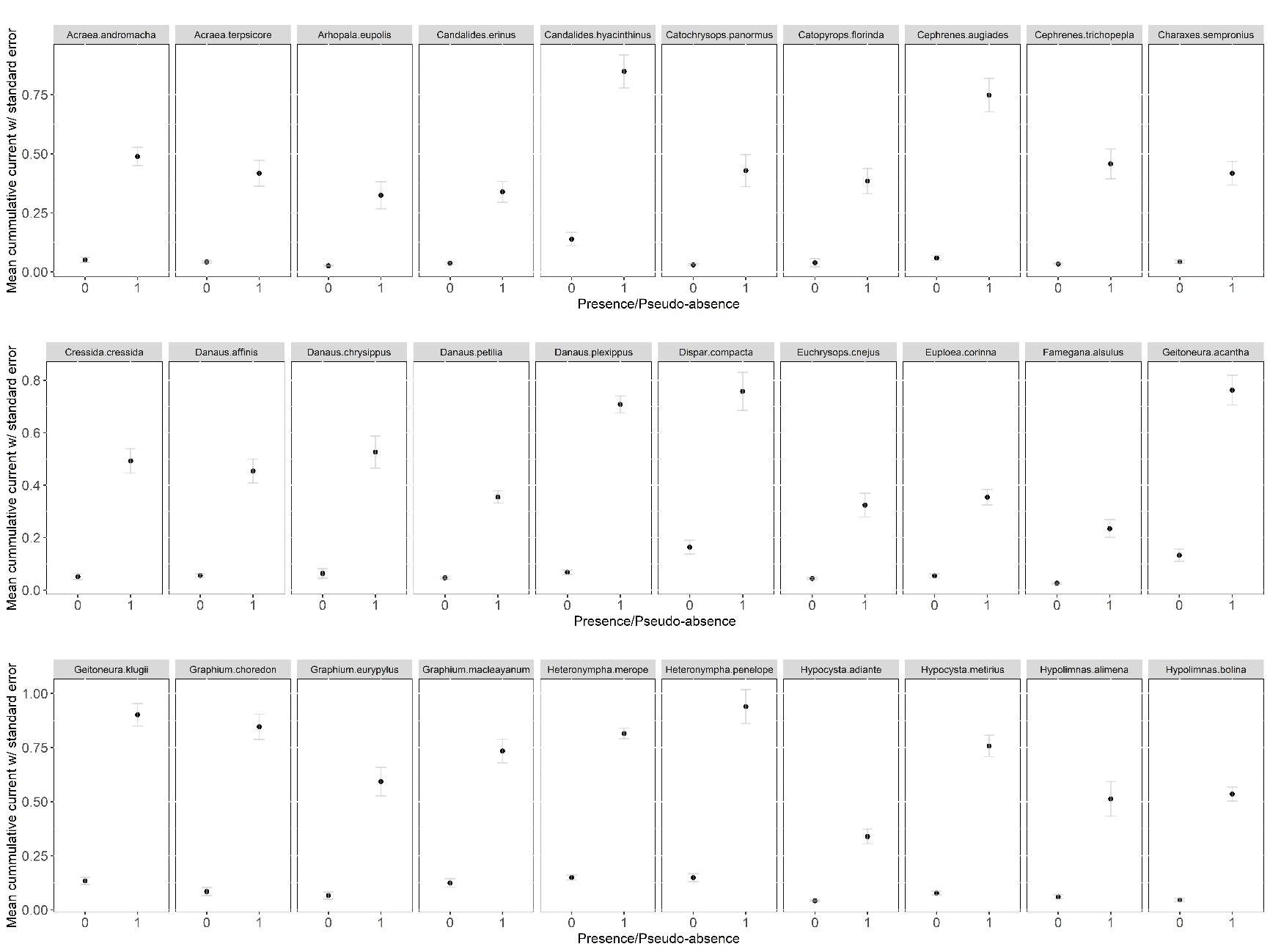

**Fig A1.** *Mean cumulative current of the presence points (1) and pseudo-absence (0) of the* functional connectivity models *of different species.*

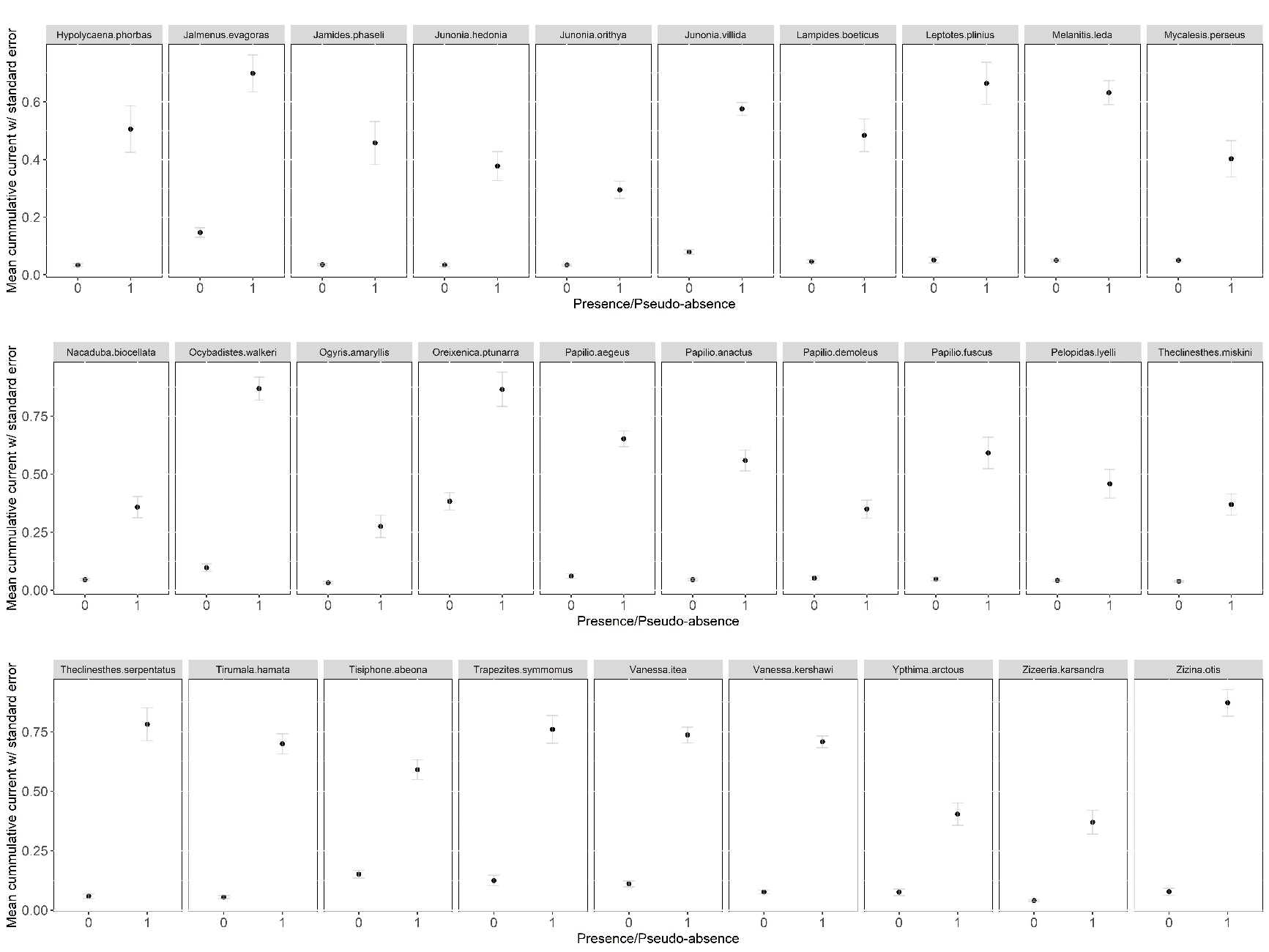

**Fig A2.** Mean cumulative current of the presence points (1) and pseudo-absence (0) of the functional connectivity models of different species.
